# Cooled SPAD array detector for low light-dose fluorescence laser scanning microscopy

**DOI:** 10.1101/2021.08.03.454878

**Authors:** Eli Slenders, Eleonora Perego, Mauro Buttafava, Giorgio Tortarolo, Enrico Conca, Sabrina Zappone, Agnieszka Pierzynska-Mach, Federica Villa, Enrica Maria Petrini, Andrea Barberis, Alberto Tosi, Giuseppe Vicidomini

**Author notes:** These authors contributed equally (Eli Slenders, Eleonora Perego).

## Abstract

The single-photon timing and sensitivity performance and the imaging ability of asynchronous-readout single-photon avalanche diode (SPAD) array detectors have opened up enormous perspectives in fluorescence (lifetime) laser scanning microscopy (FLSM), such as super-resolution image scanning microscopy and high-information content fluorescence fluctuation spectroscopy (FFS). However, the strengths of these FLSM techniques depend on the many different characteristics of the detector, such as dark-noise, photon-detection efficiency, after-pulsing probability, and optical-cross talk, whose overall optimization is typically a trade-off between these characteristics. To mitigate this trade-off, we present a novel SPAD array detector with an active cooling system, which substantially reduces the dark-noise without significantly deteriorating any other detector characteristics. In particular, we show that lowering the temperature of the sensor to −15°C significantly improves the signal-to-noise ratio due to a 10-fold decrease in the dark-count rate compared to room temperature. As a result, for imaging, the laser power can be decreased by more than a factor of three, which is particularly beneficial for live-cell super-resolution imaging, as demonstrated in fixed and living cells expressing GFP-tagged proteins. For FFS, together with the benefit of the reduced laser power, we show that cooling the detector is necessary to remove artifacts in the correlation function, such as spurious negative correlations observed in the hot elements of the detector, i.e., elements whose dark-noise is substantially higher than the median value. Overall, this detector represents a further step towards the integration of SPAD array detectors in any FLSM system.

**SIGNIFICANCE:** Single-photon avalanche diode (SPAD) array detectors are revolutionizing fluorescence laser-scanning microscopy (FLSM). Thanks to their single-photon timing and sensitivity ability and their imaging faculty, a SPAD array detector transforms any FLSM into a super-resolution microscope, and opens a whole range of possibilities for the study of sample dynamics by means of fluorescence fluctuation spectroscopy (FFS). However, dark-noise can be a severe problem for both imaging and FFS. For imaging, the signal overcomes noise only for a relatively high illumination intensity, which can be detrimental for live-cell experiments. For FFS, the noise leads to artifacts in the correlation curves, potentially leading to wrong conclusions about the sample. We show that lowering the temperature of the detector to −15°C solves both problems

## INTRODUCTION

Fluorescence laser scanning microscopy (FLSM) is one of the most powerful experimental tools in life sciences thanks to its ability to observe (sub-)cellular structures and bio-molecular processes under physiological conditions [1, 2]. Together with many other technical aspects, the choice of the detector is crucial to exploit the full potential of FLSM, or to implement a specific FLSM technique (e.g., imaging, fluorescence fluctuation spectroscopy, and their combinations with fluorescence lifetime measurements).

For FLSM-based fluorescence fluctuation spectroscopy (FFS), single-photon avalanche diodes (SPADs) are desirable. FFS is a family of techniques able to measure the mobility of and interactions between (bio-)molecules [3]. These methods rely on measuring fluctuations in the fluorescence intensity arising from a population of bio-molecules passing through the FLSM probing (or detection) volume. Because of the low photon-fluxes typically observed in these experiments - the fluorescence signal is generated from only a few fluorophores at a time - the single-photon sensitivity of SPADs is very important. Furthermore, SPADs allow recording the single-photon signal with a temporal precision (photon-timing precision) of tens of picoseconds with respect to a reference event - usually the pulsed laser excitation event; thus these detectors are ideal for time-correlated-single-photon counting (TCSPC), which is the basis for fluorescence lifetime measurements and other time-resolved experiments. For FLSM-based imaging, photo-multiplier tubes (PMTs) are preferable to SPAD detectors because of the higher dynamic range [4]. After recording a photon, the SPADs are blind for a fixed period of time (typically, 20-200 ns), the so-called hold-off time or dead time. Hence, SPADs have a maximum photon-count rate limited by the inverse of the hold-off time (5-50 MHz) [5]. Furthermore, the detector response becomes non-linear at photon-fluxes already much lower than the maximum photon-count rate. On the other hand, although PMTs offer a superior linearity at high photon-fluxes, they are not ideal neither for FFS nor for fluorescence lifetime applications. Indeed, PMTs have a lower photon sensitivity than SPADs and the digitization of the analogue PMT output, required for the TCSPC recording, may introduce unwanted fluctuations in the signal, and lead to a worse photon-timing precision compared to SPADs [6].

Because of the growing interest in FLSM systems that can do both imaging and FFS, the last years have shown the development of new detectors which combine the best characteristics of PMTs and SPADs. Hybrid detectors (HyDs) have been designed to achieve a high dynamic-range, high photon-sensitivity, and good photon-timing precision. In the context of FFS, the high dynamic-range supports experiments with a wide range of (bio-)molecular concentrations [7]. However, the complexity needed to achieve these specifications and the use of a photocathode (as for PMTs) reduces the robustness of the HyD detectors with respect to solid-states devices, such as SPADs: SPADs are immune to magnetic fields, highly resistant to mechanical shocks, and do not suffer from “burn-in” by incident light saturation. Silicon photomultipliers (SiPMs), also called multi-pixel photon counters (MPPCs), are solid-state alternatives with a high dynamic-range. A SiPM is an array of microcells, each having a SPAD and a quenching resistor, characterized by a single analog output obtained by summing the signals of all pixels. The large number of elements in the array allows many photons to be detected simultaneously without saturation [8]. Since a SiPM is an asynchronous read-out detector (i.e., it provides an analog current every time one or more SPADs get triggered), it can also be used for photon counting and for TCSPC experiments, providing a proper signal elaboration. The elaboration is easy for photon counting but more complex for TCSPC. Indeed, it is necessary to temporally separate the signals from two or more photons arriving quasi-simultaneously. This task can be achieved efficiently by combining SiPMs with the HyD technology [9]. Even though SiPM is a pixelated detector, it provides, similar to HyD, SPAD, and PMT detectors, a single output: the spatial information of where photons hit the sensor’s active area (in the case of SiPM, which SPAD element of the array) is lost. However, it has recently been shown that having access to this spatial information can shine new light on FLSM. Indeed, by using an array of detectors to image the probing volume of the FLSM, it is possible to reconstruct super-resolved images, via the image scanning microscopy (ISM) concept [10–14]. Furthermore, when the detector array also provides a high temporal resolution (> MHz), the combined spatiotemporal information can be used to augment the information content provided by a single FFS experiment [15, 16], i.e., the so-called comprehensive correlation analysis (CCA). The same spatiotemporal information combined with a similar correlation analysis was used to implement nanoscopy imaging [17]. Finally, by combining the spatial and timing information, super-resolution fluorescence lifetime imaging [14] and nanoscopy techniques based on photon-coincidences (quantum microscopy) become feasible [18].

Asynchronous read-out SPAD array detectors solve the problem of SiPM, while maintaining the best features of all other FLSM detectors [19]. Similarly to SiPM, these detectors are composed of an array of SPADs, but each SPAD can be read independently from the others. The possibility to implement such a read-out scheme is a consequence of the fact that imaging the probing region of a FLSM requires only a few tens of SPADs (e.g., 5×5 up to 7×7), and not a million of elements as for wide-field microscopy imaging. It is worth noting that SPAD array cameras with a much higher number of elements are well established [20]. However, to efficiently transfer the enormous amount of spatiotemporal information, they typically implement synchronous read-out (i.e., frame-by-frame) and they implement the photon counting or the TCSPC features on the detector itself, thus reducing the versatility of the system [20]. Because each SPAD element provides a single digital pulse each time a photon is registered, the asynchronous read-out SPAD array maintains all the characteristics of single-element SPAD detectors, such as a high-sensitivity, optimal photon-timing precision and no read-out noise. Furthermore, as for SiPM, the multiple element architecture improves the dynamic-range.

However, to effectively revolutionise FLSM, asynchronous read-out SPAD array detectors need to display excellent performance also on other important characteristics, such as photon detection efficiency (PDE), optical cross-talk probability, after-pulsing probability, and dark-noise. For SPAD arrays, the PDE depends both on the quantum efficiency of the SPAD and the fill-factor of the array (i.e., the ratio of its sensitive area to its total area). Typically, the higher the fill-factor, the higher the cross-talk probability (i.e., the probability of an element to register a photon-event as a consequence of a photon-event in an adjacent element). The use of micro-lenses is therefore a desirable alternative to increase the fill-factor. In parallel, we have recently shown that the quantum efficiency of a SPAD array can be improved by using the Bipolar-CMOS-DMOS (BCD) fabrication technology instead of the more traditional CMOS technology [21, 22]. Indeed, BCD SPAD detectors offer superior quantum efficiency, a low after-pulsing probability, excellent photon-timing precision and a low cross-talk. These features come at the price of a higher dark-count rate (DCR, i.e., the rate of spurious avalanche events due to carriers generated within the detector in the absence of light) and a higher probability of “hot-pixels” (i.e., elements whose DCR is significantly higher than the average of the array) [22]. The DCR in a SPAD array detector can be decreased by reducing the active area of the SPAD, but at the cost of a reduced fill-factor. An alternative approach with negligible detrimental effects is reducing the operating temperature of the SPAD: cooling down silicon SPADs reduces the DCR by about a decade for every 20°C of temperature reduction [23].

In this work, we implemented an active cooling system integrated in our BCD SPAD array detector. We demonstrated that operating the cooling system at −15°C reduces the DCR by more than one order of magnitude, with a negligible increase of the after-pulsing probability, and no effect on the cross-talk probability. We then integrated the cooled SPAD array detector in a FLSM system, and we show the substantial enhancement obtained both in the context of imaging and FFS. We performed imaging of fixed and live-cells, and we investigated the diffusion of freely diffusing beads in aqueous suspension and GFP-tagged proteins in living cells. These experiments demonstrated that the cooled detector allows (i) reducing the illumination power, or equivalently the pixel dwell-time, to achieve a high signal-to-noise ratio and the resolution predicted by image-scanning-microscopy (ISM) and (ii) removing the artifacts introduced in FFS experiments by the dark-noise, thus obtaining sound and robust molecule mobility information. This issue makes ISM among the most gentle super-resolution techniques for live-cell investigations. We believe that this work represents an important step towards the integration of SPAD array detectors in any FLSM and, more in general, a significant contribution to the so-called single-photon revolution: fluorescence photons from the sample are collected one-by-one with a series of signatures (e.g., in time and space), whose analysis can generate new insights about the sample, otherwise lost by conventional light recording.

## MATERIALS AND METHODS

### Cooling system

To reduce the DCR of the 5×5 BCD SPAD array detector described in [22], and thus to fully exploit the PDE enhancement (with respect to the CMOS-based counterpart [22]), we designed a compact electro-mechanical system capable of effectively cooling the device, while keeping it compatible with the existing fluorescence laser scanning microscopy setup. A two-pieces machined aluminum chamber was designed to hermetically isolate the device from the environment, thus avoiding water condensation on cold surfaces and minimizing the heat exchange (Fig. 1). The SPAD array chip was directly mounted onto a 2-stage thermo-electric cooler (TEC) having a maximum heat transfer power of 1.1 W. The TEC hot side dissipates heat through the baseplate of the chamber. The temperature sensing was done with a negative temperature coefficient (NTC) resistor, and the cooling was controlled with a dedicated control circuit. Electrical connections were routed from the chip to an embedded printed circuit board (PCB) through 25 μm bonding wires, and finally reached the acquisition system via a high-density flat-flex cable. The chamber baseplate and cover were sealed using a vacuum-grade o-ring, and the internal volume was filled with Argon gas to further reduce the heat exchange between the surfaces. Fluorescence photons can reach the sensor through an anti-reflection coated glass window (WG11010-A, ThorLabs). The dimensions of the assembly are 50 mm × 50 mm × 23 mm.

**Figure 1:**
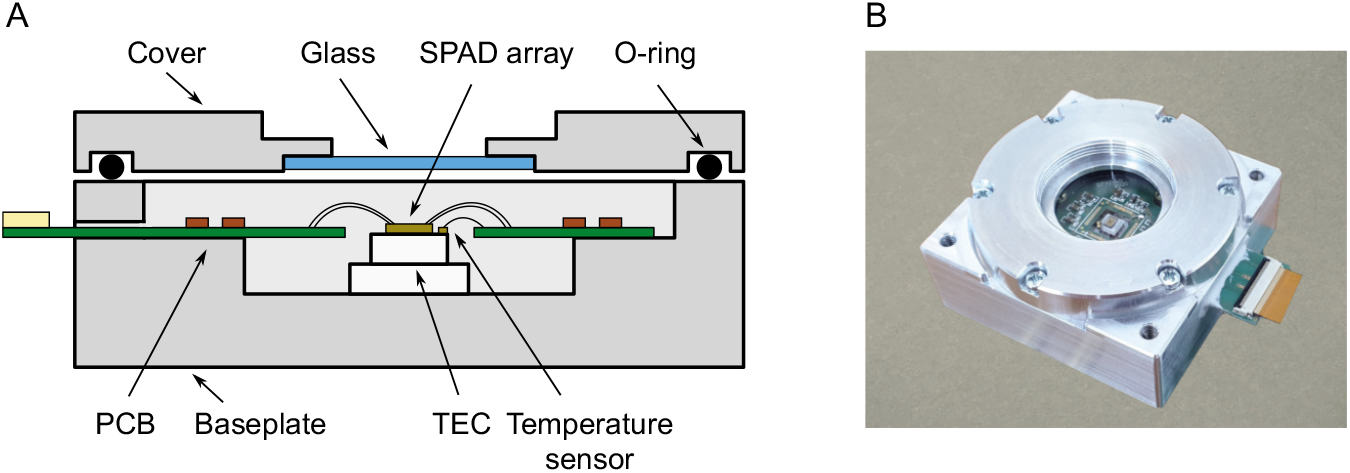
Cooled 5×5 SPAD array sensor. (A) Simplified cross-section of the 5×5 SPAD array assembly, mounted into the cooling aluminum chamber. The sensor chip was directly glued on a 2-stage TEC, floating from surrounding objects. Operating temperature is measured using an NTC resistor. Electrical connections to the chip were implemented through 25 μm bonding wires, thus minimizing heat transfer, and then transferred to an internal PCB and to a high-density flat-flex connector. The entire chamber was hermetically sealed and filled with Argon gas, to avoid condensation and to further decrease heat exchange. Photons can reach the sensor through a 1-inch AR-coated glass window. (B) Picture of the assembled sensor.

Operating temperatures ranging from room temperature down to −15°C can be chosen, while the sensor bias voltage must be adjusted accordingly to maintain a constant excess bias voltage of 5V [22]. For the data presented here, the detector hold-off time (dead time) was set to 100 ns, which is a good trade-off between a low afterpulsing probability and a high maximum count-rate.

### Microscope

The microscopy setup used both for imaging and FFS in this work is similar to the setup extensively described in [16], except for the new detector assembly. In short, a 485 nm 80 MHz pulsed laser is reflected by a dichroic beam splitter and galvanometric scanning mirrors, then sent through a Leica scan lens/tube lens system and finally focused onto the sample by a 100×/1.4 Leica oil immersion objective. The fluorescence signal is collected in de-scanned mode, passing through the dichroic beam splitter, a 488 nm notch filter and a BP500-550 emission filter. The fluorescence is finally focused onto the detector photosensitive area, with a back-projected size at the sample plane of 1.5 Airy units. We measured all the laser power values at the back-aperture of the objective lens.

We controlled the microscope with the BrightEyes control and data acquisition module (BrightEyes-CDAQM), a home-built LabVIEW (National Instruments) program based on the Carma application [24, 25]. The BrightEyes-CDAQM uses an FPGA-based data-acquisition (DAQ) card (NI USB-7856R, National Instruments), which guarantees fast prototyping and great flexibility. In short the BrightEyes-CDAQM (i) provides a graphic user interface to control the major acquisition parameters (e.g., scanned region, pixel-size, pixel-dwell time, (x,y) coordinates for the FCS measurement); (ii) registers (in photon-counting mode) the 25 digital signals of the detector array in temporal bins of minimum 500 ns, and in synchronization with the beam scanning system and other devices, e.g., laser shutters; (iii) visualises the recorded signals (e.g., intensity images and time-traces). A quite unique feature of the BrightEyes-CDAQM is the possibility to record, for each pixel, the fluorescence signal over multiple temporal bins, resulting in a four-dimensional photon counting (intensity) data structures *I*(*t*, *x*, *y*, *c*), where *t* is the time course within the pixel-dwell time, (*x*, *y*) are the spatial scanning coordinates, and *c* is the element/channel of the SPAD array detector. For FFS, the software records a photon-counting time series/trace for each SPAD array channel, i.e., *I*(*t*, *c*), where *t* are the 500 ns temporal bins. The BrightEyes-CDAQM stores both FFS and image data in HDF5 files.

### Samples

#### Fixed beads

Yellow-green carboxylate fluoSpheres (REF F8803, 2% solids, 100 nm diameter, exc./em. 505/515 nm, Invitrogen, Thermo Fisher Scientific) were diluted 1000× in ultrapure water and sonicated for 10 minutes in a water bath sonicator (Labsonic LBS1-0.6, FALC Instruments). 100 *μL* of the suspension was poured onto a cover slip previously coated with a poly-L-lysine solution (0.01%). After 8 minutes, the residue was removed and the cover slip was mounted on a microscope slide with a mounting buffer.

#### Beads in suspension

Yellow-green carboxylate fluoSpheres (REF F8787, 2% solids, 20 nm diameter, actual size 27 nm, exc./em. 505/515 nm, Invitrogen, Thermo Fisher Scientific) were diluted 5000× in ultrapure water at room temperature. Before each measurement, the bead solution was sonicated for 10 minutes in a water bath sonicator (Labsonic LBS1-0.6, FALC Instruments). A droplet of the bead solution was poured onto a cover slip for the FFS measurements.

#### Fixed cells

Hela cells were cultured in Dulbecco’s Modified Eagle Medium (Gibco™, ThermoFisher Scientific) supplemented with 10% fetal bovine serum (Sigma-Aldrich, Steinheim, Germany) and 1% penicillin/streptomycin (Sigma-Aldrich) at 37 °C in 5% CO_2_. One day before immunostaining, the cells were seeded onto coverslips in a 12-well plate (Corning Inc., Corning NY, USA). Cells were incubated in a solution of 0.3% Triton X-100 (Sigma-Aldrich) and 0.1% glutaraldehyde (Sigma-Aldrich) in the BRB80 buffer (80 mM Pipes, 1 mM EGTA, 4 mM MgCl, pH 6.8, Sigma-Aldrich) for 1 min. Hela cells were fixed with a solution of 4% paraformaldehyde (Sigma-Aldrich) and 4% sucrose (Sigma-Aldrich) in the BRB80 buffer for 10 min and then washed three times for 15 min in phosphate-buffered saline (PBS, Gibco™, ThermoFisher). Next, cells were treated for 10 min with a solution of 0.25% Triton-X100 in blocking buffer (solution of 3% bovine serum albumin (BSA, Sigma-Aldrich) in BRB80 buffer), and washed three times for 15 min in PBS. After 1 h in blocking buffer, Hela cells were incubated with monoclonal mouse anti-*α*-tubulin antibody (Sigma-Aldrich) diluted in the blocking buffer (1:1000) for 1 h at room temperature. The anti-*α*-tubulin antibody was revealed with anti-mouse Alexa Fluor 488 (Invitrogen, ThermoFisher Scientific). Hela cells were rinsed three times in PBS for 15 min. Finally, the cover slips were mounted onto microscope slides (Avantor, VWR International, Milano, Italy) with ProLong Diamond Antifade Mountant (Invitrogen, ThermoFisher Scientific).

#### Live-cells

HEK293T cells were cultured in DMEM (Dulbecco’s Modified Eagle Medium, Gibco™, ThermoFisher Scientific) supplemented with 1% MEM (Eagle’s minimum essential medium), Non-essential Amino Acid Solution (Sigma-Aldrich), 10% fetal bovine serum (Sigma-Aldrich) and 1% penicillin/streptomycin (Sigma-Aldrich) at 37 °C in 5% CO_2_. HEK293T cells were seeded onto a *μ*-Slide 8 Well plate (Ibidi GmbH, Gräfelfing, Germany). HEK293T cells were co-transfected with (SEP)-tagged-*β3* subunit GABAA receptor, a pH-sensitive variant of GFP, [26, 27] and *α*1 subunit GABAA receptor [28]to form functional *α β* GABAA receptors, for the live-cell time-lapse experiment and with pcDNA3.1(+)eGFP (Addgene plasmid #129020) for the FFS measurements. Transfection was performed using Lipofectamine™ 3000 Transfection Reagent (Invitrogen, ThermoFisher Scientific) according to manufacturer’s protocol. U2-OS cells stably expressing eGFP-DEK fusion protein were cultured in McCoy’s 5a modified medium (ThermoFisher Scientific) supplemented with 10% fetal bovine serum (Sigma-Aldrich, Germany) and 1% penicillin/streptomycin (Sigma-Aldrich, Germany) at 37°C in 5% CO_2_. Cells were seeded on a *μ*-Slide 8-Well plate Glass Bottom (Ibidi GmbH, Germany, n. 80827) 48h before the measurement. Measurements were performed in Live Cell Imaging Solution (ThermoFisher Scientific) at room temperature.

### Imaging

#### Fixed beads

We recorded bead images of 512×512 pixels, with a pixel size of 39 nm and a pixel dwell time of 100 *μ*s. We split each data-set into two time bins of 50 *μ*s each. All photons arriving during the first 50 *μ*s were used to build a first image, and likewise for the second image. We used the two images to evaluate the effective resolution by means of the Fourier ring correlation (FRC) analysis.

#### Fixed cells

We recorded cell images of 1500×1500 pixels, with a pixel size of 50 nm and a pixel dwell time of 50 *μ*s, divided into 100 time bins of 500 ns each. Images were taken with the SPAD array detector at room temperature and at −15°C.

#### Live-cells

We recorded live-cell images of 1000×1000 pixels with a pixel size of 55 nm and a dwell time of 30 *μ*s per pixel, at room temperature. The laser power was 12 *μ*W. One single image was acquired approximately every 3 minutes, for a total time of 1 hour of total acquisition. We reconstructed all the super-resolved images as described in the section below.

### Image reconstruction and analysis

We processed the imaging data-set with the Miplib library in Python [29, 30]. The Miplib library contains the adaptive pixel reassignment method for ISM reconstruction [14] and the Fourier ring correlation analysis for the evaluation of the image effective spatial resolution [29, 31]. The FRC analysis typically requires two “identical” images, which we obtained by using the ability of the BrightEyes-CDAQM to temporally split the fluorescence signal of each pixel in different time-windows ’ within the pixel dwell-time. To compare the resolution as a function of the excitation laser power, we divided the pixel dwell-time into two identical time windows, yielding the intensity datasets, *I*^1^(*x*, *y*, *c*) and *I*^2^(*x*, *y*, *c*). For each dataset, we reconstructed the relative Sum 5×5 images and adaptive pixel reassignment ISM images, which can be used for the FRC analysis.

To compare the resolution as a function of the pixel dwell time, we measured for each pixel the fluorescence signal over a time interval of 50 *μ*s, divided in time bins of 500 ns each, yielding 100 images of the sample. These were combined in post-processing to generate pairs of images with increasing pixel dwell times. For each dwell time M*t* between 500 ns and 25 *μ*s (in steps of 500 ns), we generated two independent images, *I*^1^(*x*, *y*, *c*, Δ*t*) and *I*^2^(*x*, *y*, *c*, Δ*t*) by summing the first *N* = Δ*t*/500 *ns* odd numbered and even numbered images of the time series, respectively. For each pair *I*^1^ and *I*^2^, we calculated the Sum 5 × 5 and the ISM reconstructed images and we applied the FRC algorithm.

### Fluorescence fluctuation spectroscopy

#### Beads in suspension

For beads measurements, a droplet of a freshly made bead suspension was poured onto a cover slip. Five measurements of at least 300 s each were performed with a laser power of 6.4 *μ*W, for a detector temperature of −15°C and at room temperature. The acquired time traces of the different configurations (such as *I*_12_(*t*), *I*_*Sum*3×3_(*t*), *I*_*Sum*5×5_(*t*), etc.) were split into chunks of 10 s, and the auto-correlation of each chunk was calculated using the Multipletau Python package [12]. Afterwards, all chunks from each time trace were averaged. Correlations were calculated for logarithmically scaled lag times ranging from 500 ns to 0.1 s. For scanning fluorescence correlation spectroscopy (FCS), the fluorescence time trace was recorded while scanning the laser beam in circles of 0.5 *μ*m in diameter with a frequency of 320 *μ*s per circle, by using the galvanometric scanning mirrors. For a scan speed that is fast (compared to the sample dynamics), scanning-FCS allows extracting both the beam waist and the diffusion coefficient from a single measurement [32], Supplementary Note 1. The technique can thus be considered a calibration-free alternative of conventional FCS.

#### Live-cells

HEK293T or U2-OS eGFP-DEK cells were placed on the microscope stage in a *μ*-Slide 8 Well plate with Live Cell Imaging Solution. The BrightEyes-CDAQM allows imaging of a sample, after which the user can choose the positions for FFS measurements by simply clicking on the image. Multiple FFS measurements of at least 90 s each, in different cells and in different positions within each cell, were performed with a laser power of 12 *μ*W, for a detector temperature of −15°C and at room temperature. The acquired intensity time traces of the different configurations (*I_c_*(*t*) with *c* the SPAD array element, *I*_*Sum*3×3_(*t*), *I*_*Sum*5×5_(*t*)) were split into chunks of 10 s, and the auto-correlation of each chunk was calculated using the Multipletau Python package [12], as for the beads measurements. Afterwards, all chunks from each time trace were averaged, discarding time traces that showed clear photo-bleaching. The measurements performed on the eGFP-expressing cells and in the cytoplasm of U2-OS eGFP-DEK expressing cells were fitted with a 1 component FCS model, while for the measurements on the eGFP-DEK in the nuclei, a model with anomalous diffusion was considered (see Supplementary Note 1). The least squares optimiser of the SciPy library was used to fit all the correlation curves.

## RESULTS AND DISCUSSION

### Detector characterization

We first compared the main characteristics of the SPAD array detector as a function of the temperature by implementing a series of experiments on a test-bench. To evaluate the reduction of the DCR as a function of the temperature, we placed the detector in a light-tight box and we measured the number of counts per unit of time. In the “hottest” SPAD element, the DCR decreases more than ten times: from 18.8 kHz at 25°C to 1.6 kHz at −15°C. In the central element (i.e., element 12), which is the element that typically receives the most photons in any imaging or FFS experiments, the DCR decreases more than twenty times: from 2.2 kHz to 0.1 kHz (Fig. 2 a). Despite the substantial reduction, a value of 0.1 kHz is still higher than expected: indeed, we expected a reduction of two orders of magnitude with a temperature change of −40°C. This apparent deviation is a consequence of the cross-talk effect: we repeated the experiment at −15°C by sequentially keeping only one element on at a time and found that the DCR for the central element decreases to 0.01 KHz, thus confirming the expected reduction of two orders of magnitude (Fig. 2 b).

**Figure 2:**
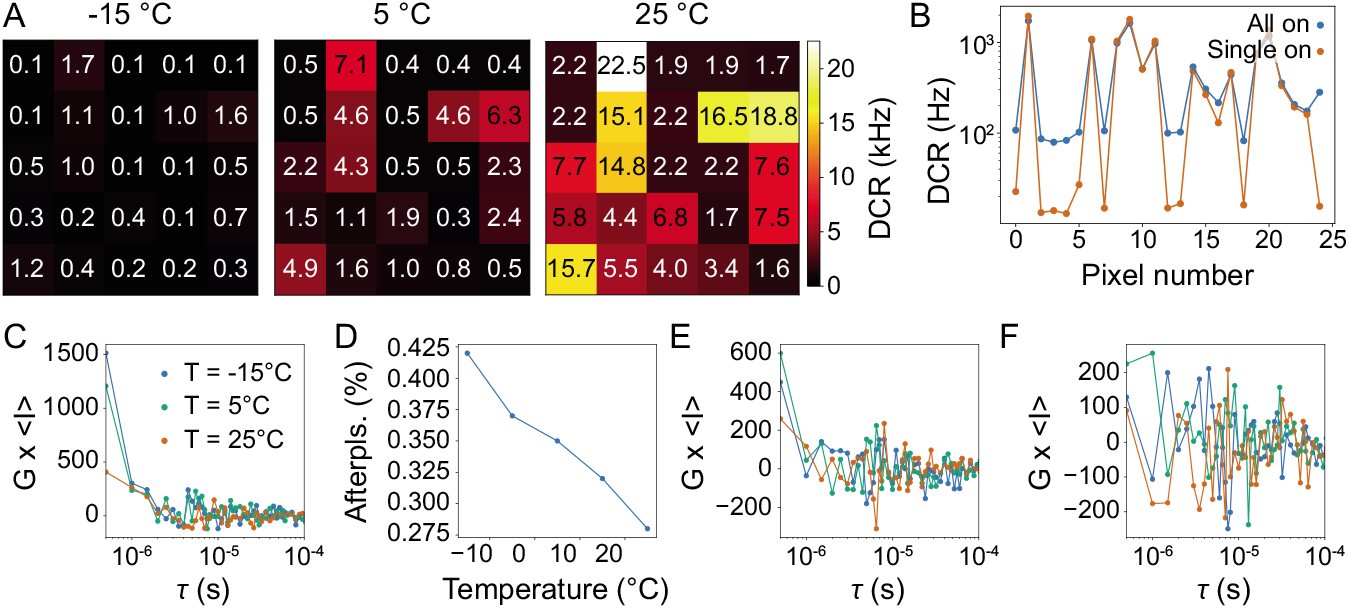
Specifications of the SPAD array sensor. (A) Dark count rate (DCR) for each pixel for various detector temperatures. The DCR was measured with all pixels simultaneously turned on. (B) Comparison between the DCR measured with all pixels simultaneously turned on (All on) and all pixels sequentially turned on (Single on) for a detector temperature of −15°C. (C) Normalized autocorrelation of the central pixel under non-correlating ambient light for different detector temperatures. (D) Afterpulsing probability as a function of the detector temperature for the central pixel. (E-F) Averaged and normalized cross-correlations between the central pixel and the four orthogonal nearest neighbours (E) or the four diagonal nearest neighbours (F) under ambient light. Same colors as in (C).

It is well known that reducing the temperature of the SPAD array can increase the after-pulsing probability [33]: the main source for after-pulses are trapped charges that are released over time, and the lifetime of the traps typically becomes longer for lower temperatures. To monitor the effect of the temperature on the after-pulsing probability, we recorded with the BrightEyes-CDAQM the intensity time-traces produced by the detector when illuminated uniformly by an uncorrelated source, such as a light-emitting diode (LED). We then calculated the auto-correlation functions multiplied by the average intensity (in the range 2-35 kHz) to take into consideration the DCR variation [34]. The curves clearly show a correlation at short lag times (< *μ*s), which are induced by the after-pulsing effect (Fig. 2 c). Since the BrightEyes-CDAQM has a time-resolution of 500 ns, we implemented the same experiment with a different data-acquisition system to quantify the increase in after-pulsing probability as a function of the temperature [22]. In particular, we used a time-correlated-single-photon counting card to measure the inter-arrival times between consecutive output pulses of an individual pixel. The contribution of the DCR to the inter-arrival times histogram can be fitted with an exponential decay at long inter-arrival times and then subtracted from the experimental data in order to have only the contribution of avalanches from the after-pulses. The after-pulse probability is then computed as the integral sum of the after-pulsing events, divided by the integral sum of the histogram itself (i.e., the total number of avalanches) (Fig. 2 d). By reducing the temperature from 25°C down to −15°C, the after-pulsing probability increases from 0.27% to 0.42%, but remains relatively low, being fully compatible with FFS.

To quantify the effect of the temperature on the optical cross-talk between the SPAD array elements, we used the same setup with the BrighEyes-CDAQM as for the after-pulsing characterization. However, instead of calculating the auto-correlation, we calculated the cross-correlation between adjacent elements. Neither the correlation curves between orthogonal neighbors, nor between diagonal neighbors, show substantial changes as a function of the temperature (Fig. 2 e,f), thus demonstrating that by cooling the detector the optical cross-talk probability does not degrade.

In short, cooling the BCD SPAD array to −15°C drastically decreases the DCR without substantially worsening any other characteristics, such as the after-pulsing probability and the optical cross-talk. Notably, by cooling this detector to −15°C, the DCR performance becomes similar to our CMOS SPAD array detector [14, 16, 29], which is, to the best of our knowledge, the only SPAD array detector effectively used so far to implement both ISM and CCA. On top of this, our BCD SPAD array offers superior characteristics in terms of PDE and after-pulsing probability [22].

### Imaging

To demonstrate the benefits of the cooled SPAD array detector in terms of imaging, we first used a calibration sample composed of 100 nm fluorescent beads. We compared imaging of beads acquired with the detector running at room temperature (uncooled) and at −15°C (cooled) for a series of different excitation laser powers (Fig. 3). For a relatively high laser power (920 nW), cooling the detector has a relatively small effect on the image quality. The contribution of the dark noise to the overall fluorescent signal is low. As a result, the images made with the cooled and uncooled detector look very similar and the FRC analysis (Fig. 3 b-d) reveals only a minor improvement in the ISM resolution upon cooling. However, for lower laser powers, the image quality with the uncooled detector deteriorates quickly. With 70 nW, the beads hardly protrude above the noise level and the FRC resolution drops by 53%; from about 200 nm at 920 nW to more than 300 nm at 70 nW. Lowering the laser power also negatively affects the resolution of images taken with a cooled detector, but in this case the difference is only 21%; from about 192 nm at 920 nW to 233 nm at 70 nW. Cooling the detector allowed lowering the laser power by more than a factor of 3 without sacrificing the resolution (Fig. 3 d). This result clearly illustrates the importance of cooling the detector for long-term live-cell imaging, in which phototoxicity and photobleaching effects come into play.

**Figure 3:**
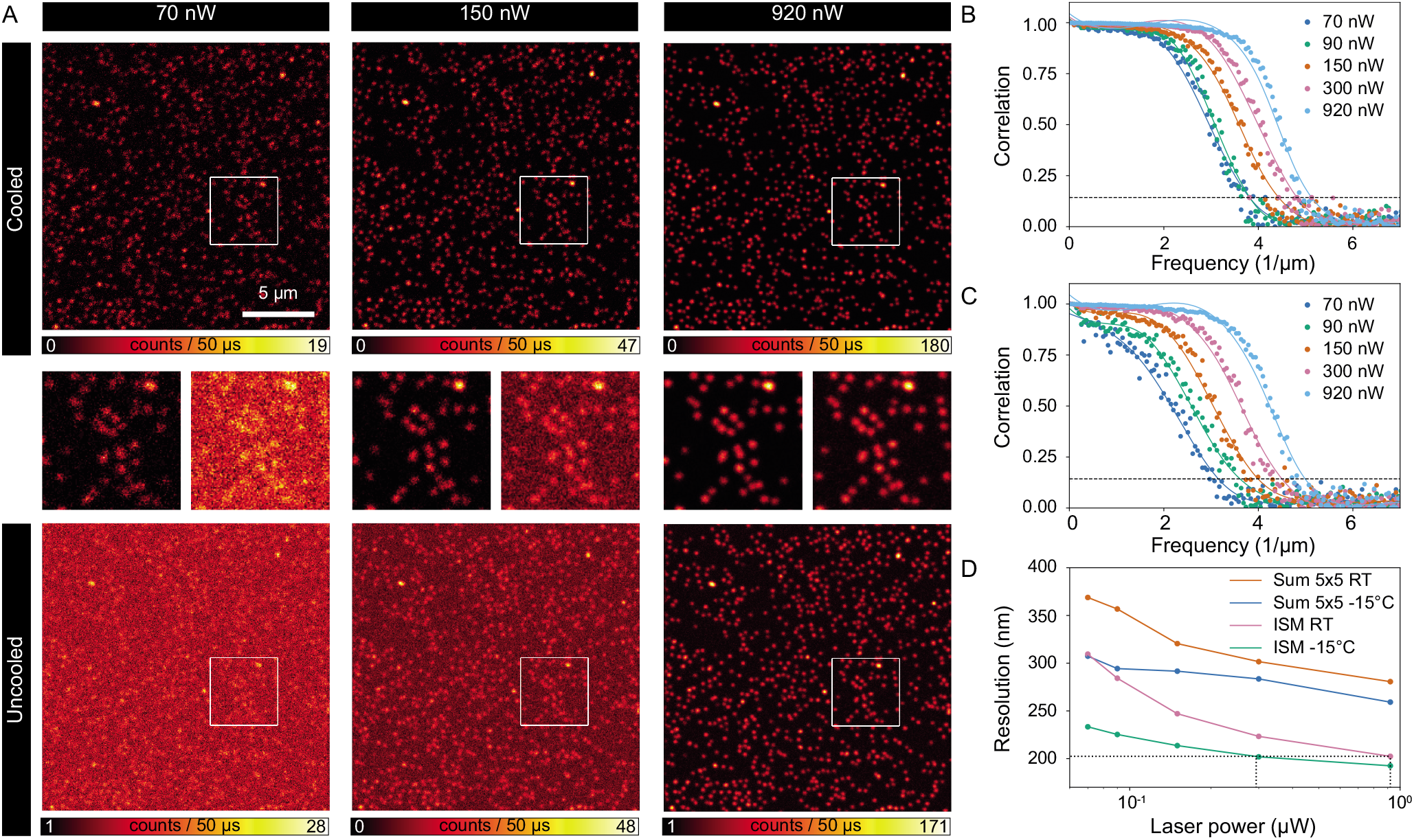
Comparison imaging fixed beads with a cooled (−15°C) vs. uncooled (25°C) SPAD array detector. Sample: 100 nm yellow-green beads. Laser powers were measured at the back focal plane of the objective and are average powers of an 80 MHz pulsed laser. (A) Example of some of the ISM images. The indicated square regions of length 4.7 *μ*m are shown enlarged in the middle row (left: cooled; right: uncooled). (B, C) FRC curves for cooled (B) and uncooled (C) imaging for various laser powers. The corresponding values for the resolution (D) were calculated from the intersection of the FRC curves with the indicated horizontal line at *y* = 1/7. (D) FRC resolution as a function of the laser power for the cooled (−15°C) and uncooled (RT) detector. Curves for both confocal imaging (Sum 5×5) and ISM are plotted. The same ISM resolution at RT with a laser power of 920 nW is reached with a cooled detector with only 294 nW (linear interpolation), black lines.

Alternatively, a cooled detector can be used to either decrease the laser power or to increase the imaging speed while maintaining the signal-to-noise ratio (SNR) and thus the resolution obtained with an uncooled detector. To validate this claim, we compared the FRC-based resolution between cooled and uncooled detector imaging of fixed Hela cell as a function of the pixel-dwell time (Fig. 4). We performed the comparison both for confocal and ISM imaging. As expected, for all the imaging conditions tested (i.e., confocal, ISM, cooled, and uncooled) the resolution improves for increasing pixel-dwell times. Notably, the ISM resolution at room-temperature (25°C) decreases faster than the confocal microscopy resolution at −15°C. Consequently, for long enough pixel-dwell times, i.e., high enough SNRs, ISM at RT performs better than confocal microscopy with a cooled detector. However, when comparing confocal microscopy with a cooled or uncooled detector, or ISM with a cooled or uncooled detector, it is clear that cooling the detector allows reducing the pixel-dwell time by more than a factor of two while maintaining the resolution obtained with an uncooled detector.

**Figure 4:**
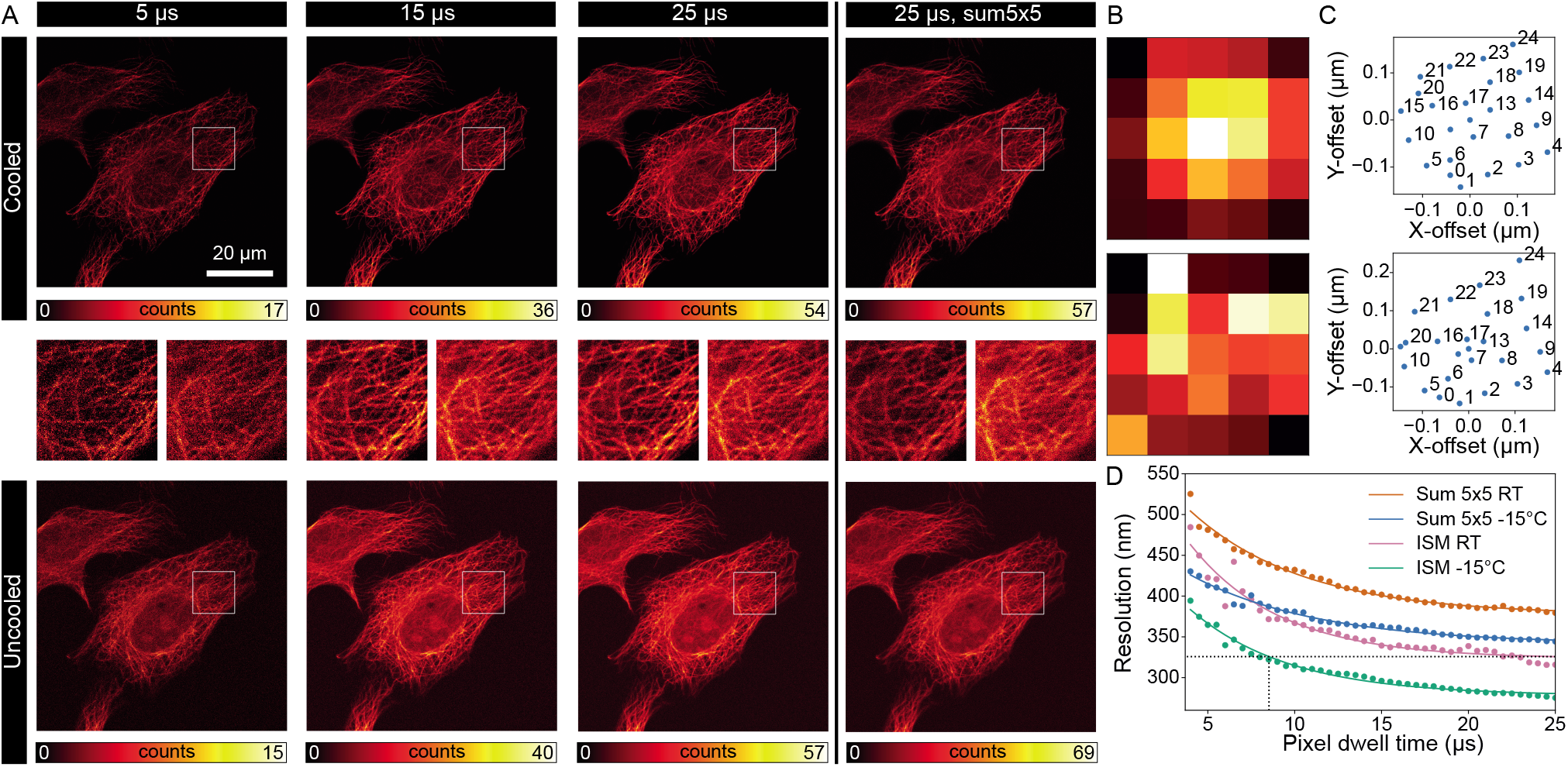
Comparison cooled vs. uncooled SPAD array detector for different pixel dwell times in images of fixed HeLa cells with alpha-tubulin staining. Laser power 100 nW at the back focal plane of the objective. The SPAD array detector was cooled down to −15°C or used at room temperature (RT/uncooled), respectively. (A) ISM images for various pixel dwell times and comparison with the Sum 5×5 image for a pixel dwell time of 25 *μs*. The middle row shows zoomed-in images of the indicated square regions of length 12.5 *μ*m of the top row images (left) and the bottom row images (right). (B) Fingerprint patterns averaged over the entire image for the images taken with a pixel dwell of 25 *μs* with the cooled (top) and uncooled (bottom) detector and (C) the corresponding shift vectors for the pixel reassignment in ISM. Some data points are not labelled for visualization purposes. (D) FRC-based resolution as a function of the pixel dwell time calculated from two independent images of the cell sample under the same experimental conditions. The continuous lines are single-exponential fits. The same FRC resolution of 326 nm can be obtained with a pixel dwell time of 25 *μs* and the detector at RT and with a pixel dwell time of 8.5 *μs* with a cooled detector (dotted line).

Finally, we used the cooled SPAD array detector on living cells, performing a time-lapse movie of cells co-expressing with the (SEP)-tagged-*β*3 and the *α*1 subunit of the GABAA receptor that co-assemble to form functional pentameric GABAA receptors expressed at the membrane surface. These receptors are known to mediate the main source of synaptic inhibition in the central nervous system. Despite the relative low laser power (12 *μ*W), we obtained high contrast and high SNR images by cooling the detector. We imaged the same cell for more than 1 hour, and we did not observe any sign of photo-toxicity or photo-bleaching (Fig. S3 and Suppl. Movie S1).

### Fluorescence fluctuation spectroscopy

To demonstrate the benefits of our new cooled SPAD array detector in the context of FFS, we performed spot-variation (or diffusion law) FCS, a technique in which the transit-time is measured as a function of the detection volume size. In particular, we compared the results of spot-variation fluorescence correlation spectroscopy (FCS) experiments performed at different detector temperatures (Fig. 5). In spot-variation FFS, the autocorrelation curves are analyzed as a function of the focal (or detection) volume size to extract information about the (bio-)molecular dynamics. In conventional spot-variation FCS, the detection volume can be adjusted by changing the radius of the confocal pinhole. In our SPAD-array based implementation, we can obtain three different volumes in a single experiment by summing - in post-processing - the signals of multiple elements of the array, i.e., *I*_12_(*t*), *I*_*Sum*3×3_(*t*), and *I*_*Sum*5×5_(*t*) intensity time-traces [15, 16]. To obtain the absolute sizes of the different detection volumes, which is needed to reveal the diffusion modality, one can use a reference sample with freely diffusing particles with a well-known diffusion coefficient. From the diffusion/transit time, it is then possible to derive the volume size, typically the lateral 1/*e*^2^ radius *ω* of a detection volume approximated by a 3D Gaussian. A different and more straightforward approach is scanning-FCS with a fast (circularly) scanning laser beam on the sample of interest. In this work, we used the second approach. We performed scanning-FCS on a solution of fluorescent beads (20 nm in diameter) diluted in water, and we calculated the autocorrelation curves for the tree different detection volumes and for two different temperature (Fig. 5 a), respectively −15°C (cooled) and 25°C (uncooled, or room temperature). By fitting the correlation curves, we extract simultaneously the transit times and the detection volume sizes (Suppl. Note 1). For the measurements at room temperature (Fig. 5 a, bottom), the amplitudes of the correlation curves differ approximately by a factor of 10 for the central pixel and 20 for Sum 3×3 and Sum 5×5 - compared to the measurements with the cooled detector (Fig. 5 a, top). Since the amplitude is inversely proportional to the number of particles in the detection volume, this discrepancy leads to inconclusive results regarding the sample concentration when the detector is employed at room temperature. Moreover, at room temperature, the correlation curves decrease at short lag times, below about 100 *μ*s, for bigger focal volumes (Sum 3×3 and Sum 5×5). This decrease, due to spurious negative correlations at short time scales in some of the pixels, is also visible in the correlation functions for a single-point acquisition FCS measurement with the uncooled detector (Fig. 5 c, left). We speculate that the decrease in the correlation is strictly connected with the dark noise signal. Indeed, (i) the correlation artifact vanishes when cooling the detector to −15°C (Fig. 5 b, left, Suppl. Fig. S2); (ii), the artifact is more evident for relatively low fluorescence intensities, i.e., low SNRs. For example, in the single-point FCS experiment, the farther the SPAD array element from the center is, the lower the fluorescence signal is and the more negative the auto-correlation becomes (Suppl. Fig. S2).

**Figure 5:**
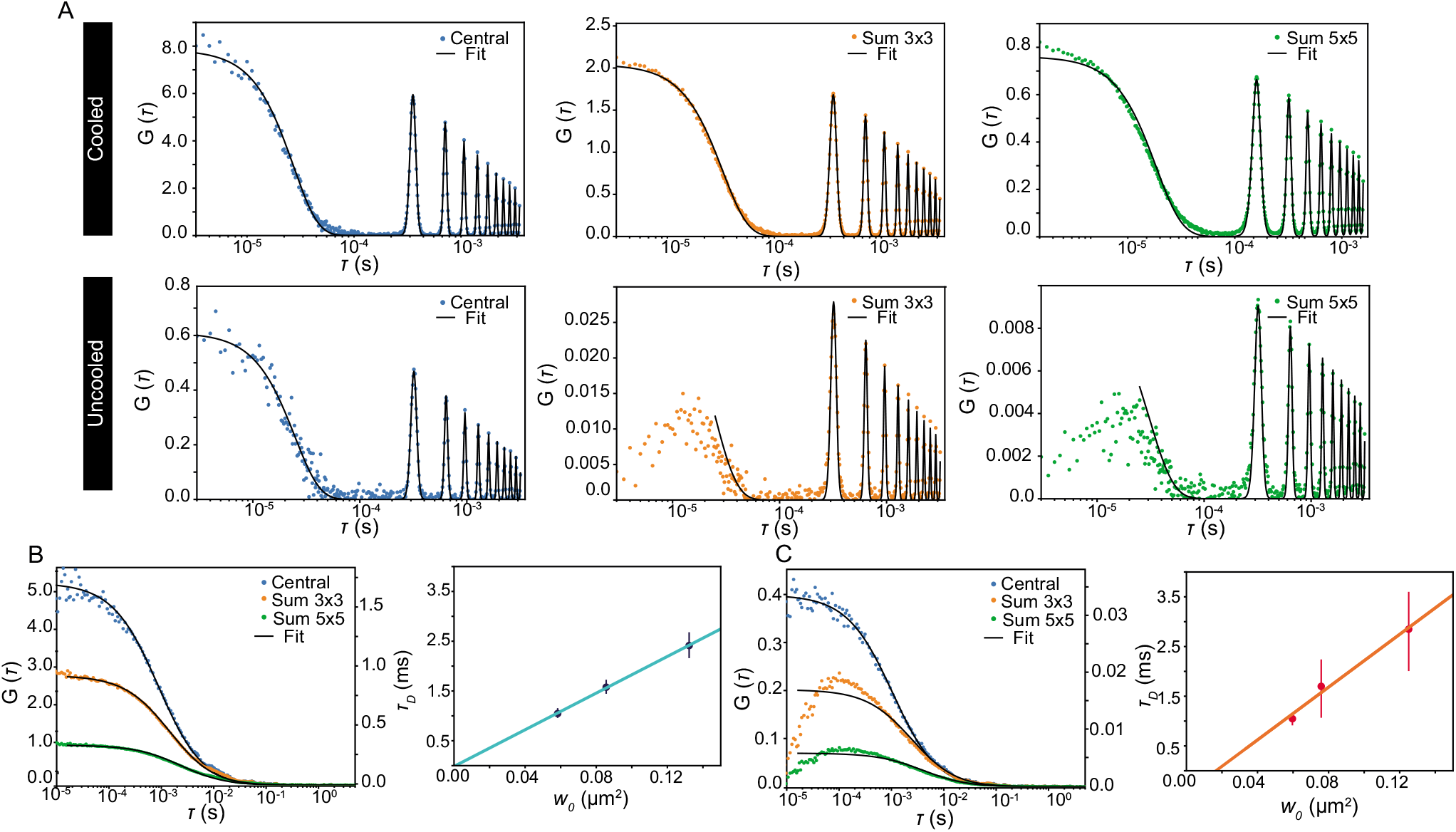
FFS measurements on fluorescent beads with a cooled (−15°C) vs. RT (25°C) SPAD array detector. Sample: freely diffusing fluorescent beads (diameter 20 nm). (A) Autocorrelation functions for a circular scanning measurement for the central pixel (left), Sum 3×3 (middle), and Sum 5 x 5 (right), for cooled (−15°C) and RT (25°C) detector. (B) Autocorrelation functions (left), for a single point measurement for the central pixel, Sum 3×3 and Sum 5×5 with the cooled detector (−15°C). Note that the scale for Sum 3×3 and Sum 5×5 autocorrelation amplitudes are reported on the right *y*-axis for visualization purposes. Diffusion times as a function of the focal volumes (right). The corresponding diffusion coefficients are (14 ± 1)*μm*^2^/*s* for the central pixel, (14 ± 2)*μm*^2^/*s* for Sum 3×3 and (14 ± 2)*μm*^2^/*s* for Sum 5×5 (averages and standard deviations over 5 measurements of 200 s each). (C) Autocorrelation functions (left), for a single point measurement for the central pixel, Sum 3×3 and Sum 5×5 with the RT (25°C) detector. Note that the scale for Sum 3×3 and Sum 5×5 autocorrelation amplitudes are reported on the right *y*/axis for visualization purposes. Diffusion time as a function of the focal volume (right). The corresponding diffusion coefficients are (15 ± 2)*μm*^2^/*s* for the central pixel, (12 ± 3)*μm*^2^/*s* for Sum 3×3 and (12 ± 3)*μm*^2^/*s* for Sum 5×5 (averages and standard deviations over 5 measurements of 200 s each).

For the uncooled detector, the lack of accurate correlation values for short lag times precludes especially the study of fast diffusion processes: the correlation curves cannot simply be cropped, as is typically done to remove the after-pulsing effects. The artifacts in the autocorrelation curves at room temperature also affect the fitted diffusion times, i.e., the characteristic lag time where the autocorrelation reaches half of its amplitude, and thus the observed diffusion modality. By using the sizes of the detection volumes extracted from the scanning FCS experiment (−15°C), we plotted the diffusion law for the single-point experiments at −15°C and at room temperature (Fig. 5 b right, Fig. 5 c right) At −15°C, the measured diffusion time scales, as expected, proportionally with the size of the detection volume (i.e., the beam waist), passing through the origin (Fig. 5 b, right), indicating that the beads are freely diffusing [35]. The corresponding diffusion coefficient is (14 ± 1)*μm*^2^/*s*,which is equivalent to a diameter of (27 ± 3) nm of the beads in according with the value provided by the manufacturer (actual size: 27 nm). On the other hand, without cooling (Fig. 5 c, right), the intercept of the linear fit with the *y* axis is negative, possibly linked with the artifacts caused by the high DCR present at room temperature. Notably, for the cooled detector, the auto-correlation curves do not show strong after-pulsing effects, not even at a relatively short lag-time (500 ns, which is the minimum sampling time of the BrigthEyes-CDAQM (Fig. Suppl. S2). This result confirms the excellent performance of the BCD-SPAD array detector in terms of after-pulsing probability - also at −15°C.

To show the applicability of the cooled SPAD-array detector, we performed FFS measurements in living cells. In particular, we performed FFS measurements in two different cell types, acquired with the cooled (−15°C) SPAD-array detector (Fig. 6). First, we measured freely diffusing GFP in HEK293T cells (Fig. 6 a). To measure the type of diffusion of the fluorescent species expressed in the cells, we employed spot-variation FCS. The fluorescence fluctuations, acquired from the chosen position (represented by a white circle in Fig. 6) within the cell cytoplasm, are summed, in a post-processing stage, to create the time traces *I*_12_(*t*), *I*_*Sum*3×3_(*t*), *I*_*Sum*5×5_(*t*)) for the different detection volumes. Here, the autocorrelation functions for the three focal volumes (Fig. 6 a, middle) central, Sum 3×3 and Sum 5×5) are analyzed with a single freely diffusing component FCS model. Indeed, the diffusion law confirms, as expected, a freely diffusing fluorescent species (Fig 6 a, right), where the diffusing times scale proportionally with the focal area. Secondly, we investigated a more complex cell system. In particular, we registered the fluorescent fluctuations at different positions - within the cell cytoplasm (Fig. 6 b) and the nucleus (Fig. 6 c) - of U2-OS cells expressing eGFP-DEK fusion protein, a chromatin architectural protein [36]. In the cell cytoplasm, DEK-GFP proteins are diffusing freely. Indeed, the spot-variation analysis (Fig. 6 c) reveals the free diffusion of these proteins, as the measured diffusion times scale proportionally with the focal volume. By fitting the autocorrelation curves with a single freely diffusing component FCS model, we found a diffusion coefficient of (38 ± 14) *μm*^2^/*s* (Fig 6 b) However, when we acquired the FFS measurements in the nuclei of these cells (Fig. 6 c), eGFP-DEK is not freely diffusing anymore. The single freely diffusing component FCS model fails to fit the autocorrelation curves acquired in the nuclei, and instead, an anomalous diffusion model has to be employed (Fig. 6 c middle). In this case, the diffusion time vs. beam waist curve does not pass through the origin, but the intercept is greater than zero, confirming the anomalous mobility of eGFP-DEK in the cell nuclei.

**Figure 6:**
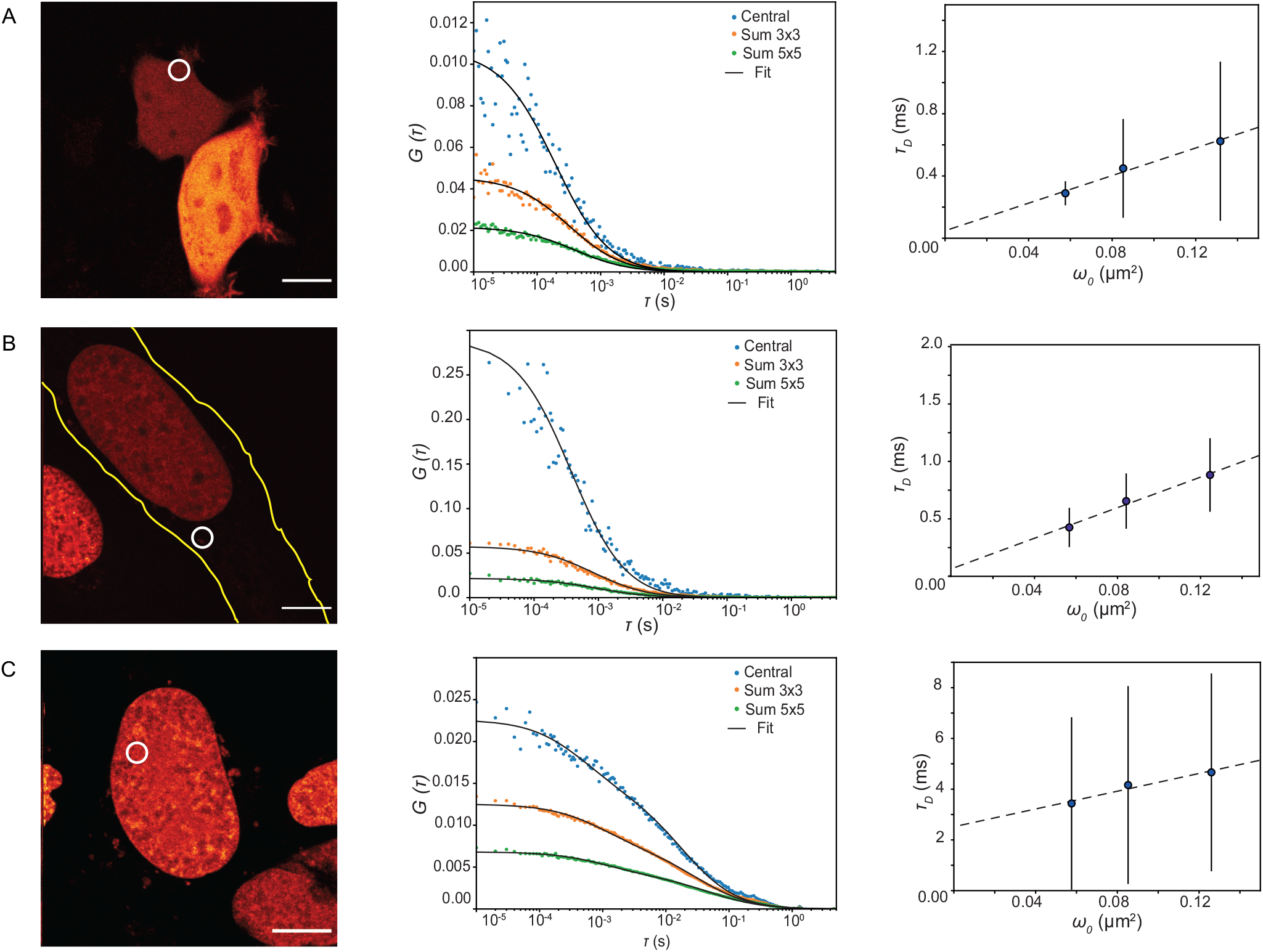
FFS measurements on live cells expressing eGFP or a GFP-tagged protein with the cooled detector. Sample: U2-OS cells expressing DEK-eGFP and HEK293T cells expressing GPF only. (A) ISM image (left) of a HEK293T cell expressing eGFP, autocorrelation curves (middle) for the central pixel, Sum 3×3, and Sum 5×5 acquired in the marked position in the cytoplasm of the cell with the cooled (−15°C) detector and diffusion time as a function of the focal volume (right). Average and standard deviations over 5 cells in 10 different positions within the cell cytoplasm. (B) ISM image (left) of a U2-OS cell expressing eGFP-DEK fusion protein (the yellow line marks approximately the cell outline), autocorrelation curves (middle) for the central pixel, Sum 3×3, and Sum 5×5 acquired in the marked position in the cytoplasm of the cell with the cooled (−15°C) detector and diffusion time as a function of the focal volume (right). Average and standard deviations over 8 cells in 18 different positions within the cytoplasm. (C) ISM image (left) of a U2-OS cell expressing eGFP-DEK, autocorrelation curves (middle) for the central pixel, Sum 3×3, and Sum 5×5 acquired in the marked position in the nucleus of the cell with the cooled (−15°C) detector and diffusion time as a function of the focal volume (right). Average and standard deviations over 5 cells in 8 different positions within the cell nuclei. The white circles mark where the FFS measurements were acquired. The scale bars are 10 *μ*m.

## CONCLUSION

Fluorescence laser scanning microscopy (FLSM) typically uses single-element detectors. Thanks to the continuous progress made on well-established technologies, such as PMTs and SPADs, and the introduction of new technologies, such HyDs and SiPMs, the performance of FLSM constantly improved and new applications emerged. However, since the 80s, it was known that FLSM imaging would benefit from a detector able to also record the spatial distribution of the fluorescence signal for each scan position (i.e., pixel), instead of integrating such a distribution across the sensitive area, as for single-element detectors. More recently, it has been demonstrated that having access to this spatial information is also important for fluorescence fluctuation spectroscopy.

Single-photon avalanche diode (SPAD) array detectors are currently the best candidate to provide this spatial information, and thus to overcome this limitation of FLSM, while maintaining the excellent characteristics of state-of-the art single-element detectors. While SPAD array detectors have been extensively optimised for specific applications of light detection and ranging (LiDAR), their usage in the context of FLSM has been a recent development, and thus a lot of room for optimization still exists. For example, the cooling system of the SPAD array to reduce the dark-noise cannot be easily implemented for the extreme conditions of many LiDAR applications, but it is fully compatible with microscopy applications.

In this work, we proposed a new cooled SPAD array detector, and we showed how cooling the SPAD array is beneficial for imaging and can even be necessary for FFS. For imaging, cooling the detector decreases the DCR, which leads to a better signal-to-noise ratio and consequently a higher contrast and better resolution. In comparison with the uncooled detector, images can be taken at almost three times the speed or a three times lower laser power for the same image quality. This is especially important for long-term live-cell imaging in which the laser powers have to be kept low to reduce phototoxicity effects and photobleaching during time-lapse experiments, and the pixel-dwell time set the frame-rate of the temporal series. This last aspect is particularly important for the combination resonant-scanner and SPAD array detector: the number of line-scans to achieve a significant signal-to-noise ratio can be reduced. For FFS, cooling the detector is necessary to remove artifacts in the correlation curves at short lag times and to get reliable results for the concentration and the diffusion time.

An underestimated characteristic of SPAD array detectors is their absence of read-out noise. Reducing the DCR of such a detector therefore effectively allows imaging affected by photon-counting noise only. Notably, single-photon megapixel CMOS camera’s have also been proposed in the context of wide-field imaging. We expect that having access to truly photon-counting imaging will open up new exciting data analysis and reconstruction methods.

As a matter of fact, the reduction of dark-noise obtained by cooling the detector, the enhancement of quantum efficiency by means of new SPAD fabrication technologies (e.g. the BCD), and, in perspective, the increased fill-factor with micro lenses and the major versatility with larger SPAD arrays (e.g., 7 × 7) are substantially reducing the motivations to not install a SPAD array detector in any FLSM.

## AUTHOR CONTRIBUTIONS

M.B., G.T., F.V., A.T., and G.V. conceived the idea. M.B., E.C., F.V., and A.T. developed the cooling system for the detector. E.S., E.P., G.T., and G.V., designed and realised the optical setup. E.S., G.T., and G.V. designed and implemented the BrightEyes-CDAQM starting from Carma. E.S. and E.P., developed the software analysis for the FFS experiments. E.S., G.T., and G.V., developed the software for image reconstruction. E.S., E.P., M.B., G.T., F.V., E.P., A.B., A.T. and G.V. designed the experiments. E.S., E.P., M.B., G.T., and E.C., characterized the SPAD array detector. E.S., E.P. and S.Z. performed the FFS and imaging experiments and analysed the data with G.V.. S.Z., A.P.-M., En.P. prepared the fixed- and live-cells. E.P., A.P.-M., S.Z. and En. P., analysed the live-cell experiments. E.S., F.V., A.B., A.T. and G.V. supervised the project. E.S., E.P. and G.V. wrote the paper with contributions from all of the authors.

## ACKNOWLEDGMENTS

The authors thank Dr. Simonluca Piazza and Dr. Marco Castello from Genoa Instruments, Dr. Paolo Bianchini and Prof. Alberto Diaspro from Istituto Italiano di Tecnologia, Ryu Nakamura from Nikon Instruments for the useful discussion about the next generation of SPAD array detectors for FLSM; Prof. Joerg Enderlein and Dr. Ingo Gregor for support in the fluorescence fluctuation spectroscopy analysis and experiments; Dr. Francesco Nicassio and Dr. Roberto Gianbruno from Istituto Italiano di Tecnologia for the pcDNA3.1(+)eGFP plasmid; Dr. Michele Oneto from Nikon Imaging Center at the IIT and Marco Scotto from Istituto Italiano di Tecnologia for the strong support in all the phases of the project; Prof. Dr. Elisa Ferrando-May from the Department of Biology, Bioimaging Center, University of Konstanz, Konstanz, Germany, for the U2-OS eGFP-DEK expressing cells; all members of the RNA Initiative at the Istituto Italiano di Tecnologia for their contribution at the long term vision of this project. This research was supported by Fondazione San Paolo, “Observation of biomolecular processes in live-cell with nanocamera”, No. EPFD0098 (E.S., S.Z., and G.V.), by the European Research Council, Bright Eyes, No. 818699 (G.T., and G.V.), and by the European Union’s Horizon 2020 research and innovation programme under the Marie Skłodowska-Curie grant agreements No 890923 (SM-SPAD) (E.S.), and No 841661 (qCHROMDEK) (A.P.-M.).

