## Supplementary notes for "Cooled SPAD array detector for low light-dose fluorescence laser scanning microscopy"

[1]Molecular Microscopy and Spectroscopy, Istituto Italiano di Tecnologia, 16152 Genoa, Italy;
[2]Dipartimento di Elettronica, Informazione e Bioingegneria, Politecnico di Milano, 20133 Milan, Italy;
[3]Nanoscopy and NIC@IIT, Istituto Italiano di Tecnologia, 16152 Genoa, Italy;
[4]Synaptic Plasticity of Inhibitory Networks, Istituto Italiano di Tecnologia, Genoa, Italy;
[5]These authors contributed equally.

An online supplement to this article can be found by visiting BJ Online at http://www.biophysj.org.

### Supplementary Note 1

In conventional FCS, in which the excitation beam is kept steady, the correlation function $G(\tau)$, for one single component, is given by [1]

$$G(\tau) = \frac{1}{N\left(1 + (\frac{\tau}{\tau_D})^\alpha\right)\sqrt{1 + (\frac{\tau}{\tau_D})^\alpha \frac{1}{SP^2}}}. \tag{1}$$

N is the average number of particles in the focal volume, $\tau_D$ the characteristic transit time of a particle diffusing through this volume, and SP the shape parameter of the point-spread-function (PSF). We approximate the PSF as a 3D Gaussian function with a lateral $1/e^2$ radius of $\omega$ and a $1/e^2$ height of $SP$ x $\omega$. Then, $\tau_D = \frac{\omega^2}{4D}$, with D the diffusion coefficient. We also include in the model for the correlation the anomalous parameter $\alpha$, which describes the degree of diffusion anomality. When $\alpha$ is 1, the species is diffusing freely.

Since the fit parameter $\tau_D$ contains both the size of the volume and the diffusion coefficient, it is impossible to extract both parameters from the same experiment. Therefore, in conventional FCS, $\omega$ is derived from a reference experiment with a sample with a known D.

In alternative, the two parameters $\omega$ and D can be decoupled by scanning the laser beam. In this case, $G(\rho, \tau)$ is a function of both spatial shifts $\rho$ and temporal lags $\tau$:

$$G(\rho, \tau) = \frac{1}{N\left(1 + \frac{\tau}{\tau_D}\right)\sqrt{1 + \frac{\tau}{\tau_D SP^2}}} \exp\left(-\frac{\rho^2}{\omega^2\left(1 + \frac{\tau}{\tau_D}\right)}\right). \tag{2}$$

In the case of a laser beam scanning in circles, Fig. S1, $\rho(\tau)$ is

$$\begin{aligned}\rho(\tau) &= \sqrt{R^2 + R^2 - 2R^2\cos(\gamma(\tau))}\\ &= \sqrt{2R^2(1 - \cos(\tfrac{2\pi}{T}\tau)},\end{aligned} \tag{3}$$

with T the time needed to scan a full circle.

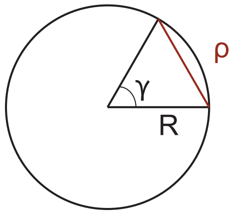


Figure S1: **Sketch circular scanning FCS.** R is the radius of the scan pattern. When the laser beam has travelled over an angle $\gamma$ in a time interval $\tau$, then $\rho(\tau)$ corresponds to the distance between these two points. Note that $\rho$ is not the distance travelled by the laser beam in this time interval.

Combining Eq. 2 and 3 yields

$$G(\rho,\tau) = \frac{1}{N\left(1+\frac{4D\tau}{\omega^2}\right)\sqrt{1+\frac{4D\tau}{\omega^2 SP^2}}} \exp\left(-\frac{4R^2\sin^2\left(\frac{\pi}{T}\tau\right)}{\omega^2+4D\tau}\right). \quad (4)$$

Since $D$ and $\omega$ now appear in two different forms in the equation, as a fraction and as a sum, both parameters are decoupled and can be fit together in a single experiment.

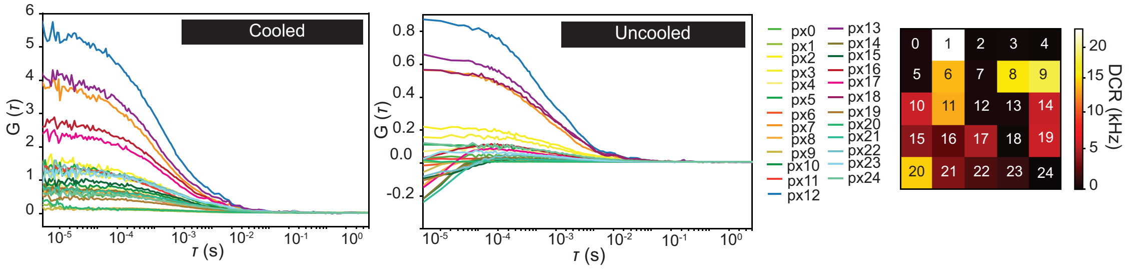


Figure S2: **Autocorrelation curves from each single detector pixels for cooled (-15°C) and uncooled (RT) detector.** Sample: fluorescent beads 20 nm of diameter. Negative correlation are visible in the autocorrelation curves acquired with the detector at RT (middle), in comparison to the autocorrelation curves for the same sample acquired at a detector temperature of -15°C (left). The negative spurious correlation at low lag times are evident especially for the detector pixels showing high DCR, as shown in the fingerprint of the detector acquired at RT (right).

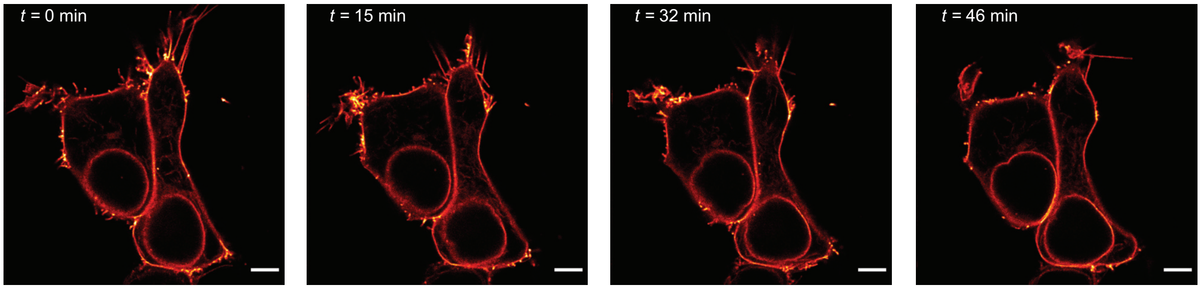


Figure S3: **Time lapse images of a living cell.** Sample: HEK cell co-transfected with (SEP)-tagged-$\beta 3$ and $\alpha 1$ subunits of the GABAA receptor forming pentameric GABAA receptors expressed at the membrane surface The images are acquired with a laser power of 12 $\mu$W with the detector at -15°C for more than 1 hour. The scale bar is 10 $\mu$m.
